# variancePartition: Interpreting drivers of variation in complex gene expression studies

**DOI:** 10.1101/040170

**Authors:** Gabriel E. Hoffman, Eric E. Schadt

## Abstract

As genomics studies become more complex and consider multiple sources of biological and technical variation, characterizing these drivers of variation becomes essential to understanding disease biology and regulatory genetics. We describe a statistical and visualization framework, variancePartition, to prioritize drivers of variation with a genome-wide summary, and identify genes that deviate from the genome-wide trend. variancePartition enables rapid interpretation of complex gene expression studies and is applicable to many genomics assays.

---

High-throughput genomics assays have revolutionized our understanding of the molecular etiology of human disease, molecular biology of cell lineage and genetic regulation of gene expression. Transcriptome profiling in particular has been widely applied to detect variation in transcript levels attributable to differences in disease state, cell type or regulatory genetics. As transcriptome profiling studies have expanded in size and scope, they have grown increasing complex and consider multiple sources of biological and technical variation. Recent studies have simultaneously considered multiple dimensions of variation to understand the impact of cell type^1^, tissue type^2^, brain region^3^, experimental stimuli^4^, time duration following stimulus^5^ or ancestry^1,4,6^ on the genetic regulation of gene expression. More studies are including a disease axis, for example to characterize the role of regulatory genetics on late onset Alzheimer’s disease in multiple brain regions^7^.

The fundamental challenge in the analysis of complex datasets is to quantify and interpret the contribution of multiple sources of variation. Indeed the most pressing questions concern the relationship between these sources of variation. How does cell or tissue type effect the genetic regulation of gene expression, and does it vary by ancestry^1,2^? What is the relative contribution of experimental stimulus versus regulatory genetics to variation in gene expression^5^? Is technical variability of RNA-Seq low enough to study regulatory genetics and disease biology, and what are the major drivers of this technical variability^2,8,9^? A rich understanding of complex datasets requires answering these questions with both a genome-wide summary and gene-level resolution.

Recently, statistical methods that decompose variation in gene expression into the variance attributable to multiple variables in the experimental design have yielded valuable insight into the biological and technical components driving expression variation^8,10–12^. Yet statistical limitations of previous methods and the lack of a convenient and scalable implementation for analysis and visualization have prevented wider application of this analysis framework.

In order to improve interpretation of these complex datasets, we have developed a statistical and visualization framework, variancePartition. variancePartition leverages the power of the linear mixed model^13^ to jointly quantify the contribution of multiple sources of variation in high-throughput genomics studies. In applications to transcriptome profiling, variance Partition fits a linear mixed model for each gene and partitions the total variance into the fraction attributable to each aspect of the study design, plus the residual variation. Because it is built on the first principles of the linear mixed model, variancePartition has well characterized theoretical properties and accurately estimates the variance fractions even for complex experimental designs where the standard ANOVA method is inadequate (Supplementary Note). Moreover, variancePartition gives strong interpretations about the drivers of expression variation, and we demonstrate that these findings are reproducible across multiple datasets.

As an example we consider the 660 RNA-Seq experiments from the GEUVADIS study^6,8^ of lymphoblastoid cell lines from 462 individuals of 5 ancestries and 2 sexes sequenced across 7 laboratories. For a single gene, the total variation can be partitioned into the contributions of these components of variance plus residual variance:

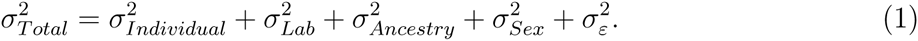

The contribution of each driver of variation can be interpreted based on the fraction of total variation it explains. Thus the fraction of variance due to variation across individuals is

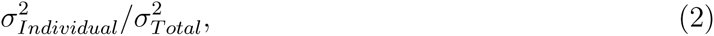

and the fractions from all components of variation sum to 1.

Because these statistics are simple to describe, the variancePartition framework can give particular insight into how each of these dimensions of variation drive transcriptional variability. A typical variancePartition analysis comprises 1) fitting a linear mixed model to quantify the contribution of each dimension of variation to each gene, 2) visualizing the results, and 3) integrating additional data about each gene to interpret drivers of this variation. Here we apply variancePartition to four gene expression studies to demonstrate how the method facilitates interpretation of drivers of expression variation in complex study designs with multiple dimensions of variation. While these datasets are already well-characterized, we illustrate how variancePartition enables rapid interpretation of complex datasets.

Applying variancePartitition to the GEUVADIS^6,8^ dataset illustrates how the method can decouple biological and technical variation, and further decompose biological variation into multiple components. Expression variation across individuals, ancestries and sexes is biological, variation across the labs where the samples were sequenced comprise technical artifacts, while the residual variation remains uncharacterized. Results from representative genes illustrate how variancePartition identifies genes where the majority of variation in expression is explained by a single variable such as individual or sex, while variation in other genes is driven by multiple variables (Figure 1a, Supplementary Table 1). Since the variance fractions sum to 1 for each gene, it is simple to compare results across genes and across sources of variation. Visualizing these results genome-wide illustrates that variation across individuals is the major source of expression variation and explains a median of 55.1% of variance genome-wide (Figure 1b). The median variance explained by laboratory (6.8%), ancestry (4.9%) and sex (0%) is substantially smaller. We note that the variance explained by individual increases to 63.8% when ancestry is removed from the analysis since ancestry is a biological property of each individual (Supplementary Figure 1, Supplementary Note).

**Figure 1:**
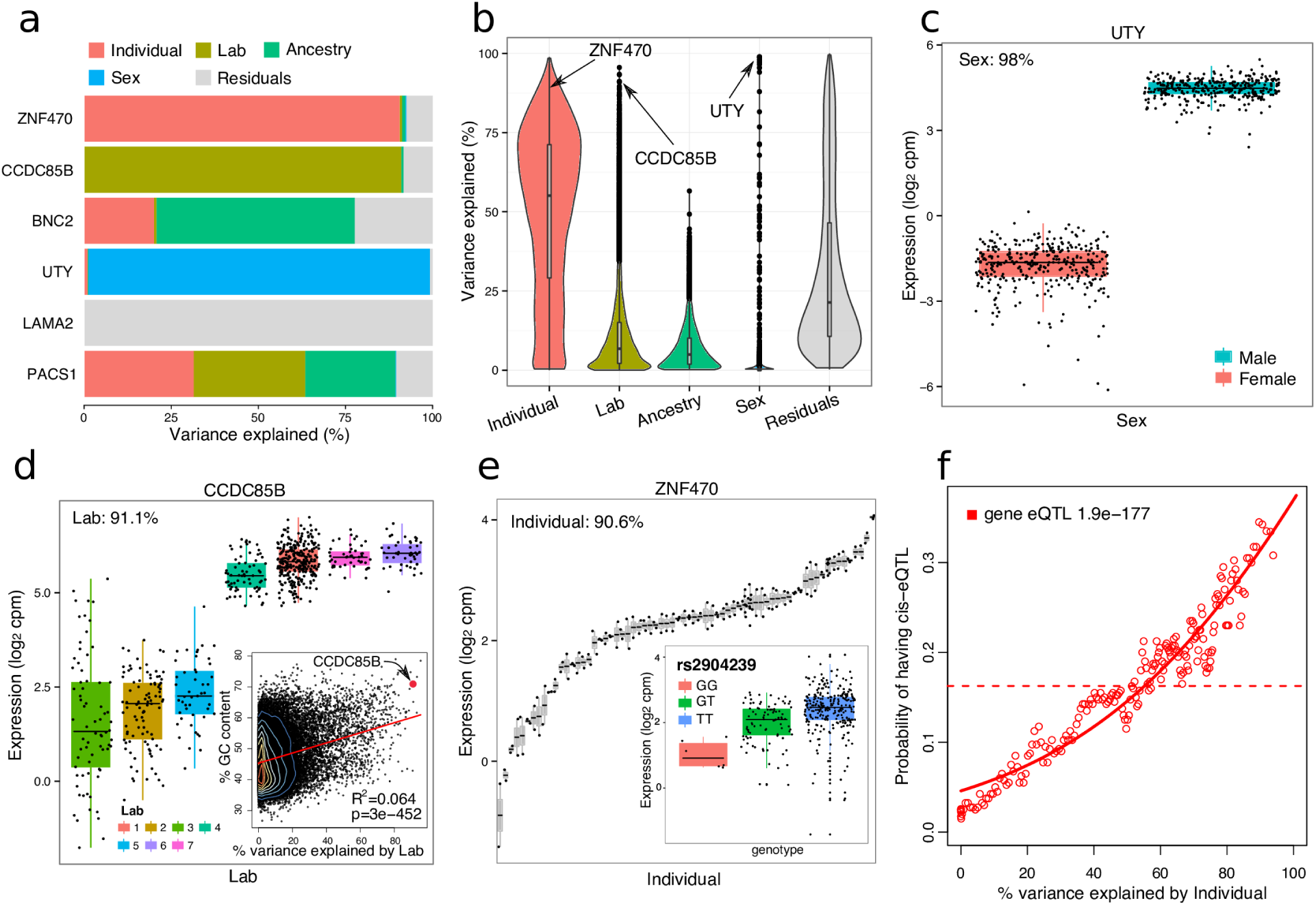
Analysis of GEUVADIS dataset identifies drivers of expression variation. (**a**) Total variance for each gene is partitioned into the fraction attributable to each dimension of variation in the study. (**b**) Violin and box plots of percent variation in gene expression explained by each variable. Three representative genes and their major sources of variation are indicated. (**c**) Boxplot of UTY expression stratified by sex. (**d**) Boxplot of CCDC85B expression stratified by lab. Inset shows scatter plot of percent GC content versus percent variance explained by lab. Red line indicates linear regression line with coefficient of determination and p-value shown. (**e**) Boxplot of ZNF470 expression stratified by individual for a subset of individuals with at least 1 technical replicate. Inset illustrates a cis-eQTL for ZNF470 where expression is stratified by genotype at rs2904239. (**f**) Probability of each gene having a cis-eQTL plotted against the percent variance explained by individual. Dashed lines indicate the genome-wide average probability (i.e. 18% of genes have a detected eQTL in this dataset), and curves indicate logistic regression smoothed probabilities as a function of the percent variance explained by individual. Points indicate a sliding window average of the probability of genes in each window having a cis-eQTL. Window size is 200 genes with an overlap of 100 genes between windows. P-values indicate the probability that an association as strong as between percent variance and eQTL probability occurs by chance according to the logistic regression smooth.

Yet particular genes show substantial deviation from the genome-wide trend. This is particularly noticeable for sex, where of the 51 genes for which sex explains more that 10% variance 46 are on the X or Y chromosomes. For example, variation across sex explains 98% variance in UTY on the Y chromosome (Figure 1c). While differential expression measures the differences in mean expression between the sexes, variancePartition measures the variance within and between each sex. This analysis indicates that variation across sexes explains very little variation genome-wide, but has a strong effect on a small number of genes.

Integrating additional data with gene-level results from variancePartition can give a clear interpretation of the drivers of variation. For example, 91.1% of variation in CCDC85B is explained by variation across laboratory. This gene has a very high GC content of 70.9% and is consistent with the genome-wide pattern where the degree of variation across laboratories is positively correlated with GC content (Figure 1d). While technical variation in RNA-Seq is known to depend on GC content^8,9^, variancePartition gives a clear illustration of how the effect of technical artifacts varies substantially across genes. Moreover, this analysis can be used to identify other correlates underlying technical issues in expression variation.

In addition, variancePartition gives a strong interpretation to genes whose expression varies across individuals by relating the gene-level results to cis-regulatory variation. For example, the fact that 90.6% of variation in ZNF470 is explained by individual suggests that this variation is driven by genetics, and, in fact, ZNF470 has a cis-eQTL (Figure 1e). This observation is also seen genome-wide, as genes with a greater fraction of variation across individuals have a significantly higher probability of having a cis-eQTL detected in this study (Figure 1f). This analysis explicitly demonstrates how expression variation across individuals is driven by cis-regulatory variation.

Application of variancePartition to *post mortem* RNA-Seq data of multiple tissues tissues from the GTEx Consortium^2^ decouples the influence of multiple biological and technical drivers of expression variation. We analyzed 489 experiments from 103 individuals in 4 tissues (blood, blood vessel, skin and adipose tissue) in order to restict the analysis to tissues with RNA-Seq data for most individuals (Supplementary Table 2). Variation across tissues is the major source of variation (median 37.4%) while the technical variables expression batch (2.9%), ischemic time (1.2%), RNA isolation batch (0.4%), and RIN (0.2%) have a moderate effect on expression variation genome-wide (Figure 2a). Variation across expression batches is correlated with GC content but to a lesser degree that other datasets (Supplementary Figure 2). Cumulatively, these technical variables explain only 4.7% of the total expression variation. Concerns about reliability of RNA-Seq data from *post mortem* samples has been raised due to the potential effects of RNA degradation following cell death^14,15^. variancePartition analysis indicates that variation in ischemic time has as relatively small effect genome-wide and the fraction of variance it explains is comparable to other technical effects, yet the effect varies substantially across genes.

**Figure 2:**
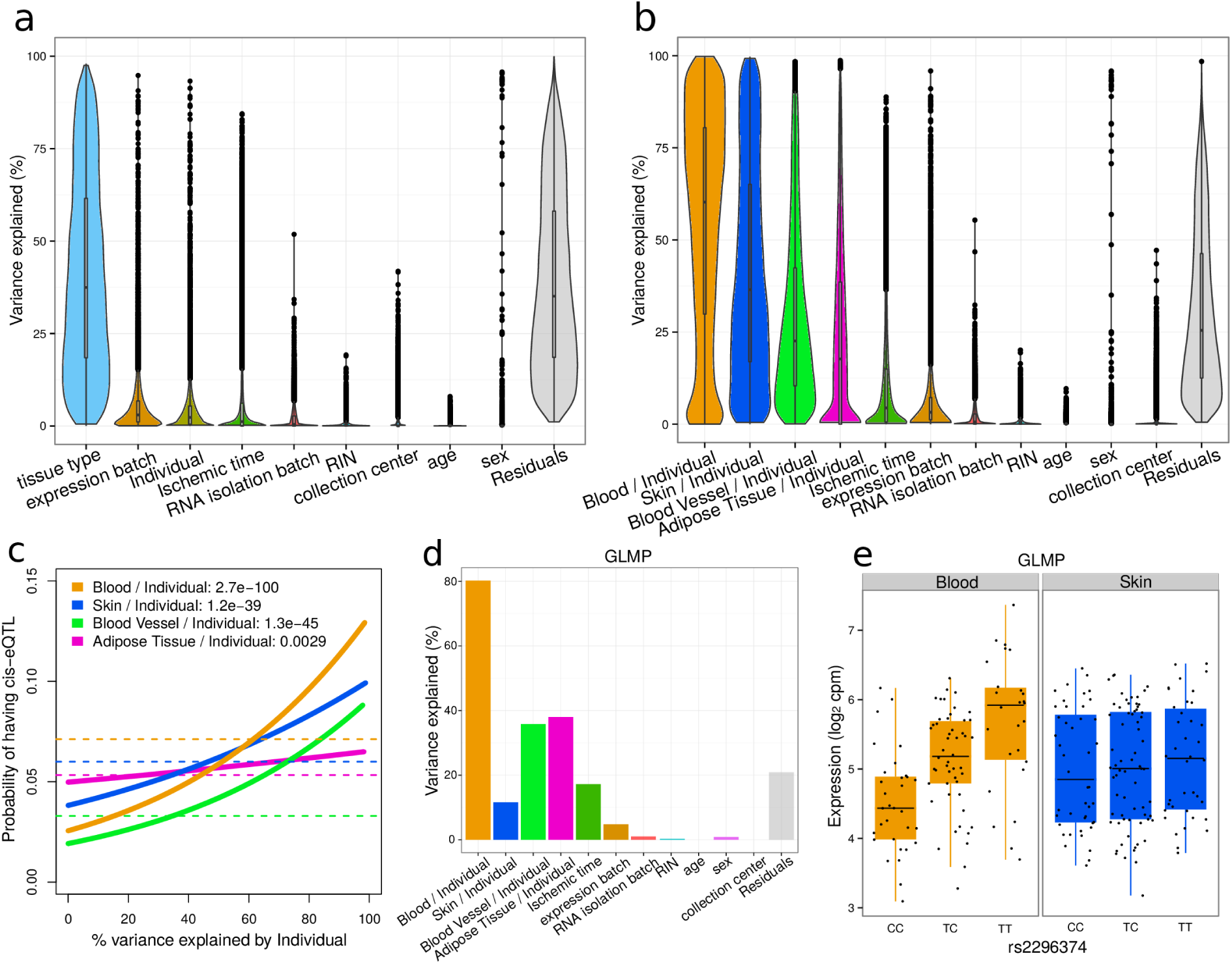
Analysis of GTEx dataset identifies drivers of expression variation at multiple levels. (**a**) Violin and box plots of percent variation in gene expression explained by each variable. (**b**) Results from variancePartition analysis allowing the contribution of individual to vary in each tissue. (**c**) Probability of each gene having a cis-eQTL plotted against the percent variance explained by individual within each tissue. Dashed lines indicate the genome-wide average probability, and curves indicate logistic regression smoothed probabilities as a function of the percent variance explained by individual within each tissue. P-values indicate the probability that an association as strong occurs by chance. (**d**) Fraction of variation in GLMP explained by each source of variation. (**e**) GLMP contains a cis-eQTL active in blood but not skin.

The flexibility of the linear mixed model framework allows variancePartition to analyze cross-individual variation within each tissue. Since the variance is analyzed within multiple subsets of the data, the total variation explained no longer sums to 1 as it does for standard application of variancePartition. Yet the results allow ranking of dimensions of variation based on genome-wide contribution to variance and enables analysis of gene-level results (Supplementary Note). While variation across individuals explains only a median of 2.3% of variation when all tissues types are considered together, there is substantial variation across individuals within each tissue separately (Figure 2b). Cross-individual variation is highest in blood (median 60.3%), while skin (36.5%), blood vessel (22.5%), and adipose tissue (17.7%) exhibit lower cross-individual variation. The fraction of variation explained by individual within each tissue is directly related to the probability of each gene having a cis-eQTL within the corresponding tissue (Figure 2c). This association is not as strong as in other datasets likely due to the smaller number of individuals and to the relatively small fraction of cross-individual variation in adipose tissue.

At the gene-level, varianceParition can prioritize genes based on multiple criteria. For examples, GLMP exhibits higher variation across individuals within blood but low variation in skin (Figure 2d). This is consistent of a tissue-specific regulatory variation, and, in fact, the cis-eQTL rs2296374 influences gene expression in blood but not in skin (Figure 2e).

The conclusions drawn here from variancePartition analysis are reproducible across multiple datasets, and we illustrate further applications to datasets from the ImmVar^1^ and SEQC projects^9^ (Supplementary Note, Supplementary Figures 3-4).

A variancePartition analysis produces a genome-wide summary but also reports gene-level results to identify genes that deviate from the genome-wide trend. The fraction of variation is an interpretable statistic which can easily be compared across genes, across drivers of variation and across datasets. Decomposing expression variation into its constituent parts facilitates prioritizing drivers of variation based on their contribution at both the genome-wide and gene-level. By comparing the influence of multiple drivers of variation, this analysis directly addresses questions about the extent to which biological signal or technical artifacts are driving variation in the data. Moreover, the fraction of variation is also interpretable in terms of intra-class correlation, a metric used to assess biological and technical reproducibility^13,16^(Supplementary Note).

Finally, variancePartition provides a general statistical and visualization framework applicable to studying drivers of variation in many types of high-throughput genomic assays including RNA-Seq (gene‐, exon‐ and isoform-level quantification, splicing efficiency), protein quantification, metabolite quantification, metagenomic assays, methylation arrays and epigenomic sequencing assays. Although we have focused here on large-scale studies, variancePartition analysis has given valuable insight into RNA-Seq datasets with as few as 20 samples. The variancePartition software is an open source R package and is freely available on Bioconductor. The software provides a user-friendly interface for analysis and visualization with extensive documentation, and will enable routine application to a range of genomics datasets.

## Acknowledgements

We would like to thank B. Kidd, B. Losic, N. Beckmann, R. Kosoy and many other colleagues at Mount Sinai and Stanford for providing valuable feedback on the software and manuscript.

## Author contributions

G.E.H conceived the statistical method, implemented the software and performed the analysis. G.E.H wrote the manuscript with E.E.S.

## Online Methods

variancePartition summarizes the contribution of each variable in terms of the fraction of variation explained (FVE). While the concept of FVE is widely applied to univariate regression by reporting the *R*^2^ value from a linear model, variancePartition extends FVE to applications with complex study designs with multiple variables of interest. The linear mixed model framework of variancePartition allows multiple dimensions of variation to be considered jointly in a single model and accommodates discrete variables with a large number of categories.

### Model specification

Each gene is analyzed separately using the linear mixed model

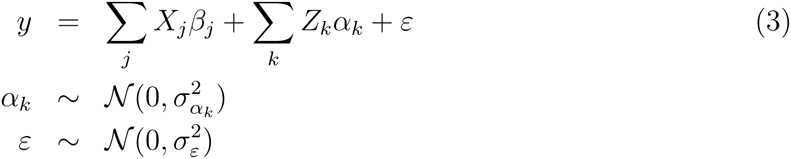

where *y* is the expression of a single gene across all samples, *X*_*j*_ is the matrix of *j*^*th*^ fixed effect with coefficients *β*_*j*_, *Z_k_* is the matrix corresponding to the *k*^*th*^ random effect with coefficients *α_k_* drawn from a normal distribution with variance 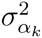. The noise term, £, is drawn from a normal distribution with variance 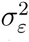. All parameters are estimated with maximum likelihood^17^ as simulations under a range of experimental designs indicate that this approach gives the most accurate FVE estimates (Supplementary Note, Supplementary Figures 5-8).

Variance terms for the fixed effects are computed using the *post hoc* calculation

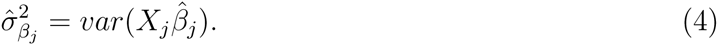

The total variance is

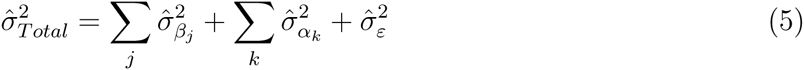

so that the fraction of variance explained by the *j*^*th*^ fixed effect is

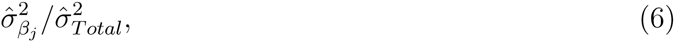

by the *k*^*th*^ random effect is

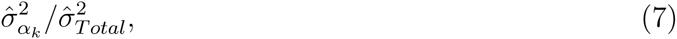

and the residual variance is

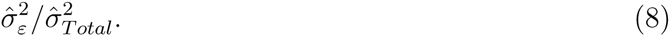

In the standard application of variancePartition, these fractions sum to 1 and are always positive by definition. Each gene is processed separately so that only visualization and reporting of genome-wide summary statistics use the results from multiple genes.

### Modeling measurement error in RNA-Seq data

Uncertainty in the measurement of RNA-Seq data can be modeled with observation-level precision weights that model the relationship between expression magnitude and sampling variance^18^. variancePartition incorporates these precision weights to create a heteroskedastic linear mixed model^17^. Denoting the vector of precision weights for a single gene across all samples as *w*, the model is fit by weighting the residual variance from Equation (3) so that

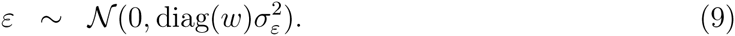

### Variation across individual within subsets of the data

The linear mixed model underlying variancePartition allows the effect of one variable to depend on the value of another variable. Statistically, this is called a varying coefficient model^13^. This analysis examines the expression variation across individuals within multiple cell types, or another subset of the data. A given sample is only from one cell type, so this analysis asks a question about a subset of the data. The data is implicitly divided into subsets base on cell type and variation explained by individual is evaluated within each subset. This subsetting means that the variance fractions no longer sum to 1, but the model still allows ranking of dimensions of variation based on genome-wide contribution to variance and enables analysis of gene-level results. See the Supplementary Note for more details.

### Smoothing eQTL results with logistic regression

A logistic regression model was used to regress the eQTL status, coded as 0 or 1, on the precent variation explained by individual. The predicted probability from this model is used to create the smoothed curves, and this probability is a monotonically increasing function of the precent variation explained by individual due to the restrictions of the logistic model. The p-values indicate a hypothesis test that the coefficient relating the eQTL probability to precent variation explained by individual is zero.

### Visualization with violin-boxplots

The fraction of variation explained by each aspect of the study design is presented using a combined violin and boxplot^19^. The boxplots indicate the median, inner quartile range (IQR) and 1.5 times the IQR. Data beyond this are plotted as points. Violin plots indicate the density of data points based on their width.

### Software

The variancePartition software v1.0.5 is available from http://bioconductor.org/packages/variancePartition/

